# TractoFlow: A robust, efficient and reproducible diffusion MRI pipeline leveraging Nextflow & Singularity

**DOI:** 10.1101/631952

**Authors:** Guillaume Theaud, Jean-Christophe Houde, Arnaud Boré, François Rheault, Felix Morency, Maxime Descoteaux

**Affiliations:** Sherbrooke Connectivity Imaging Laboratory (SCIL), Université de Sherbrooke, Canada; Imeka Solutions Inc, Sherbrooke, Canada

**Author notes:** 2500, boul. de l’Université, Sherbrooke (Québec) Canada, J1K 2R1, *Email address:* (Guillaume Theaud).

**Keywords:** Diffusion MRI, Pipeline, Processing, Tractography, Reproducibility, Nextflow, Singularity

## Abstract

A diffusion MRI (dMRI) tractography processing pipeline should be: i) reproducible in immediate test-test, ii) reproducible in time, iii) efficient and iv) easy to use. Two runs of the same processing pipeline with the same input data should give the same output today, tomorrow and in 2 years. However, processing dMRI data requires a large number of steps (20+ steps) that, at this time, may not be reproducible between runs or over time. If parameters such as the number of threads or the random number generator are not carefully set in the brain extraction, registration and fiber tracking steps, the end tractography results obtained can be far from reproducible and limit brain connectivity studies. Moreover, processing can take several hours to days of computation for a large database, even more so if the steps are running sequentially.

To handle these issues, we present *TractoFlow*, a fully automated pipeline that processes datasets from the raw diffusion weighted images (DWI) to tractography. It also outputs classical diffusion tensor imaging measures (fractional anisotropy (FA) and diffusivities) and several HARDI measures (Number of Fiber Orientation (NuFO), Apparent Fiber Density (AFD)). The pipeline requires a DWI and T1-weighted image as NIfTI files and b-values/b-vectors in FSL format. An optional reversed phase encoded b=0 image can also be used. This pipeline is based on two technologies: *Nextflow* and *Singularity*, as well as recommended pre-processing and processing steps from the dMRI community. In this work, the TractoFlow pipeline is evaluated on three databases and shown to be efficient and reproducible from 98% to 100% depending on parameter choices. For example, 105 subjects from the Human Connectome Project (HCP) were fully ran in twenty-five (25) hours to produce, for each subject, a whole-brain tractogram with 4 million streamlines. The contribution of this paper is to introduce the importance of a robust pipeline in terms of runtime and reproducibility over time. In the era of open data and open science, efficiency and reproducibility is critical in neuroimaging projects. Our TractoFlow pipeline is publicly available for academic research and is an important step forward for better structural brain connectivity mapping.

## 1. Introduction

Diffusion magnetic resonance imaging (dMRI) is the main technique to non-invasively obtain information about the white matter. Diffusion MRI is currently at the core of structural connectivity or white matter brain mapping [Van Essen et al., 2012], using dMRI tractography [Descoteaux et al., 2009; Girard et al., 2014] to reconstruct and visualize the white matter architecture [Maier-Hein et al., 2017; Jeurissen et al., 2017]. However, there could be from 20 to 25 processing steps involved to perform dMRI tractography from a raw dMRI data.

The data processing depends on many tools in different packages with many parameters. Setting up the environment and installing all dependencies to process the data can be long, tedious and difficult, even more so for a beginner in dMRI. Then, the runtime of the different tasks is variable and can take several hours. The whole processing for one subject typically takes more than 5 hours when using a pipeline that incorporates all processing steps [Cook et al., 2006; Daducci et al., 2012; Cui et al., 2013; Sotiropoulos et al., 2013]. Hence, when ran sequentially for many subjects, can take days to months.

Several users set up the environment without giving a package version number. Over time, users upgrade packages on the computer to have new functionalities or bug patches. Some tools such as virtual environments exist to create multiple working environments with proper package versionning. However, this multi-environment concept is not appropriate for the majority of non-computer scientist users.

In longitudinal studies, new datasets are often acquired months or years after running the dMRI pipeline for the first time. As just mentioned, running the same pipeline, with the same parameters and the same software versions is difficult. Moreover, several studies are based on a subset of the full database to run different experiments. After these experiments, the whole database is used to run the pipeline. In this case, it is important to run the same processes with the same parameters as the experiments. In diffusion MRI pipelines, an important argument that is often omitted is reproducibility [Roine et al., 2019].

If a diffusion MRI pipeline is ran twice on the same dataset, with the same parameters, the two results should either be identical or have a high reproducibility percentage. For tractogram reproducibility, mean that every segment of every streamlines should be the same up to the endpoint. However, algorithms are often based on random sampling for *brain extraction*, *registration* and *tractography* processing steps. For tractography algorithms, random sampling is used to set streamline starting points. Most of them set the random sampling for the whole brain. This global random sampling does not ensure the reproducibility of the seeding process: if only one voxel is removed from the seeding mask, the tractogram reproducibility could drop to 0%.

However, a small difference in a seeding mask must only have a small impact on the reproducibility of the tractogram. Registration or brain extraction algorithms are also based on random sampling. This random sampling can have a big impact on reproducibility results and must be taken into account for the future algorithms.

In the dMRI field, the ordering of processing steps (e.g. denoising, motion correction, brain extraction, etc) are different across researchers and across laboratories [Maier-Hein et al., 2017; Schilling et al., 2019]. These processing differences create an issue when sharing processed databases or to move forward to open databases. The dMRI community should have a more uniform and transparent processing to enhance the collaborations across laboratories such as the fMRI community has started to do with *fmriprep* [Esteban et al., 2018].

The efficiency of the pipeline is a keypoint for dMRI processing. To quickly process more subjects, high performance computers (HPC) are often used. However, dMRI processing pipelines are not necessarily adapted for HPC. Easy use on HPC is an important feature for the choice of the pipeline creation tool software.

In summary, problems of current dMRI pipelines are: i) not reproducible in test-test experiments because libraries and code evolve, ii) not reproducible in time, iii) often, not adapted or at least sub-optimal in terms of multi-processing and cluster usage, iv) often, not adapted or optimized for big database processing.

We introduce a new diffusion MRI tractography processing pipeline, based on Nextflow [Di Tommaso et al., 2017] and Singularity [Kurtzer et al., 2017], for human brain connectivity mapping. We call this pipeline *TractoFlow*. *TractoFlow* is publicly available for academic research purposes and code is available here. (https://github.com/scilus/tractoflow) We first describe the different datasets used in our experiments, the materials, libraries used to develop the pipeline and the advantages of using Nextflow and Singularity. Then, we enumerate the processing steps of the pipeline and the metrics used to analyze the reproducibility. Next, we present the results of the reproducibility analysis for the maps generated by the pipeline and the tractogram. We also quantify the runtime of the results. We conclude with a discussion on technology choices, current open questions and future work.

## 2. Methods

### 2.1. Datasets

The pipeline was tested on 2 different datasets created at the Université de Sherbrooke imaging center (CIMS): Penthera1.5T [Girard et al., 2015; Chamberland et al., 2019] and Penthera3T [Paquette et al., 2019]. The current study uses 30 acquisitions per dataset, 60 acquisitions in total. The study was performed according to the guidelines of the Internal Review Board of the Centre Hospitalier Universitaire de Sherbrooke.

#### Penthera1.5T

The acquisitions were done by the Sherbrooke Connectivity Imaging Lab in 2014. Ten young and healthy subjects (four women and six men, mean of 25 (+2.91) years old) were scanned four times: twice with ten minutes delay and twice with two days delay. The diffusion-weighted images (DWI) were acquired on a 1.5 Tesla MRI (Siemens, Magnetom) with a single-shot echo-planar imaging sequence (128×128 matrix, TR 11000 ms, TE 98 ms, GRAPPA factor of 2) having 64 directions distributed using b=1000 mm^2^/s, one b=0 mm^2^/s and a 2 mm isotropic spatial resolution. A reversed phase encoded b=0 image is acquired in 2 mm isotropic spatial resolution to be able to correct for EPI distortions. A T1-weighted MPRAGE image (TR 1860 ms, TE 3.54 ms) is available in 1 mm isotropic spatial resolution.

#### Penthera3T

The acquisitions were done by the Sherbrooke Connectivity Imaging Lab in 2016. Thirteen young and healthy subjects (three women and ten men, mean of 25.92 (+−1.85) years old) were scanned six times: two sessions with three scans per session. The DWI (112×112 matrix, TR 5615 ms, TE 95 ms, SENSE factor of 2) were acquired on a 3 Tesla MRI (Philips, Ingenia) with a single-shot echo-planar imaging sequence having 3 different shells, b=300, 1000, 2000 mm^2^/s with respectively 8, 32 and 60 directions distributed, 7 b=0 mm^2^/s for a total of 107 images and a 2 mm isotropic spatial resolution [Caruyer et al., 2013]. A reversed phase encoded b=0 image is available, to correct EPI distortions, with the same spatial resolution as the DWI. The T1-weighted MPRAGE image (TR 7.9 ms, TE 3.5 ms) has a 1mm isotropic spatial resolution.

#### Human Connectome Project (HCP)

A subset of the Human Connectome Projet (HCP) database [Van Essen et al., 2012] was processed with this pipeline. The 105 subjects selected are the same used in [Wasserthal et al., 2018]. The subjects are already processed with the minimal preprocessing pipelines for the HCP [Glasser et al., 2013]. The DWI were acquired with high spatial resolution of 1.25mm isotropic. The acquisition scheme contains 270 gradient directions on three b-values (90 gradient directions for each b-value) b=1000, 2000, 3000 s/mm^2^ and 18 b=0 s/mm^2^ images. The T1 weighted image is the one available in the minimal processed registered on the DWI in 1.25mm isotropic spatial resolution.

### 2.2. Materials

Bash scripts are the simplest way to create a processing pipeline but are limited. A bash script does not naturally parallelize to multiple subjects, is hard to maintain over time and does not allow to resume processing. Multiple pipeline creation tools exist and allow us to develop a pipeline more easily than with Bash script.

Our pipeline is based on 2 technologies: Nextflow [Di Tommaso et al., 2017] and Singularity [Kurtzer et al., 2017]. Nextflow is a pipeline creation tool that executes tasks based on the Dataflow paradigm. These tasks are executed in a given order and parallelized when multiple subjects are given in input. Moreover, a maximum number of CPU to use is configurable for each process in order to optimize resources use. Nextflow allows to follow a given naming convention for the output files.

Nextflow supports container software such as Singularity container (and Docker). Singularity permits storing all dependencies required by the Nextflow pipeline to execute, in a container. When the container is built, the versions of the dependencies inside are fixed and cannot be updated. This feature ensures the reproducibility of the dependencies version used.

Our Nextflow pipeline calls multiple tasks illustrated in Figure 1. A configuration file sets all of the parameters used in the different tasks (See Supplementary materials). This configuration file is adaptable. The tasks use the tools stored in the Singularity container. The tools installed in the Singularity container are FSL [Jenkinson et al., 2012], MRtrix3 [Tournier et al., 2019], ANTs [Avants et al., 2009], DIPY [Garyfallidis et al., 2014a]. The tool versions used in the Singularity container are listed in the Supplemetary materials.

**Figure 1:**
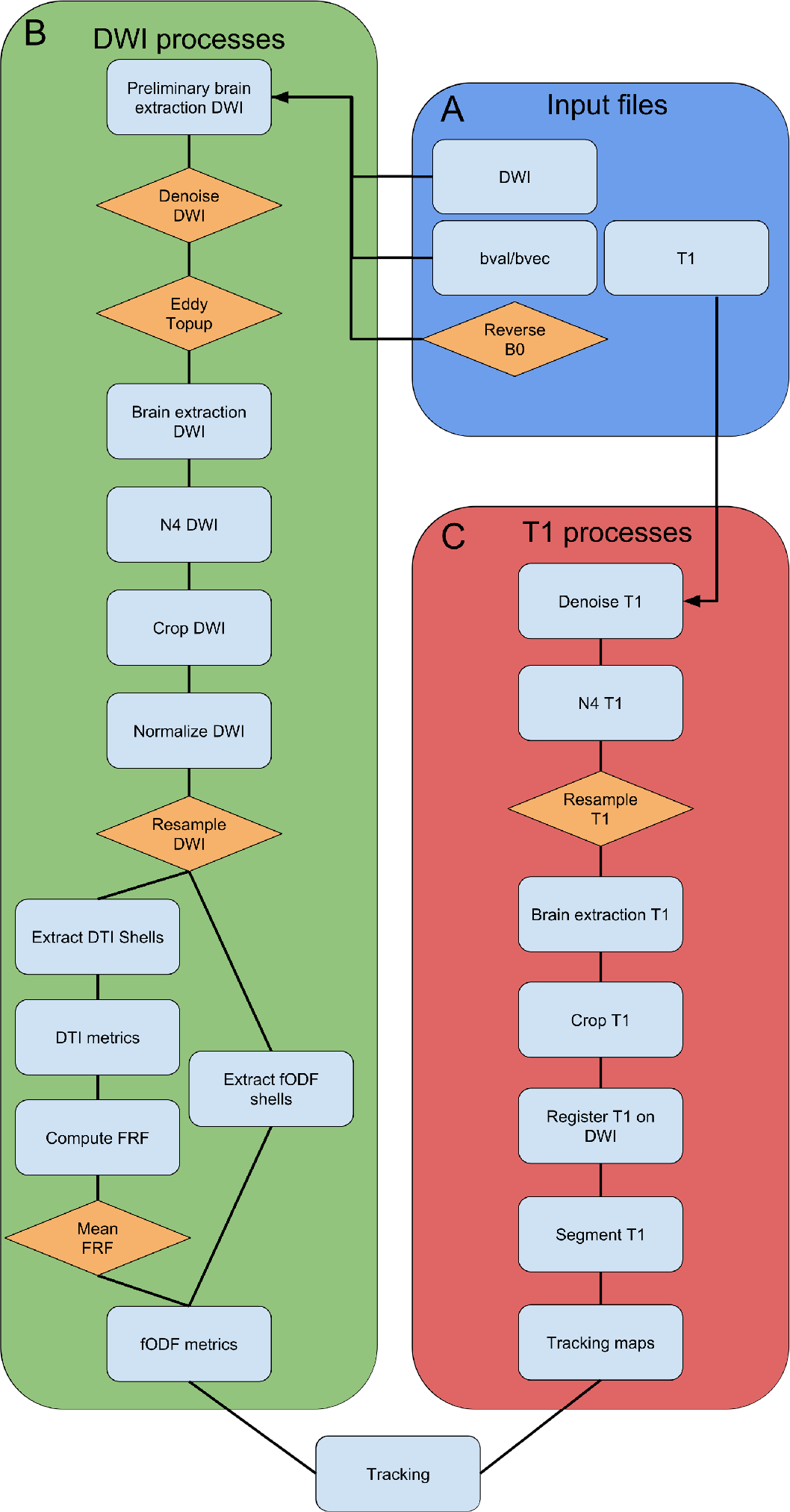
The graph of the main processes of the pipeline. In A (blue), the input files required to run the pipeline. In B (green), the DWI processes that take the DWI, the b-values/b-vectors files and the reversed phase encoded b=0 image. In C (red), the T1 processes that take the T1 weighted image in input. In orange, all processes or images that are optional.

To evaluate the reproducibility and the performances of the pipeline, we use 3 types of computers (2 clusters and 1 standard desktop), as described in Table 1.

**Table 1:** The 3 computer specifications. The CPU type (model of the processor), the architecture, the number of threads and the quantity of memory is described.

|  | Cluster 1 | Cluster 2 | Desktop |
| --- | --- | --- | --- |
| CPU type | 2 x Intel 8160F | 2 x Intel E5-2683 v4 | Intel i7-3820 |
| Architecture | Skylake | Broadwell | Sandy Bridge E |
| Number of threads | 48 | 32 | 8 |
| Memory (in Gb) | 190 | 128 | 32 |

### 2.3. Data and code availability statement

All the data and pipeline code is downloadable. The two databases, Penthera1.5T (https://doi.org/10.5281/zenodo.2602022) and Penthera3T (https://doi.org/10.5281/zenodo.2602049), are available under license. Moreover, the Singularity container (https://github.com/scilus/singularity-tractoflow) and *TractoFlow* (https://github.com/scilus/tractoflow) are available and under a license.

### 2.4. Pipeline inputs

The pipeline requires a DWI, a T1 weighted image, the b-values/b-vectors files (FSL format) and, if available, a reversed phase encoded b=0 image (See Figure 1.A) for distortion correction. The DWI, T1 weighted image and the reversed phase encoded b=0 must be in NIfTI format and the b-values/b-vectors are text files. We recommend *dicom2niix* to convert DICOM in NIfTI format. The pipeline can process multiple subjects at the same time, as long as they follow a given naming convention. Each subject must have a folder containing DWI (dwi.nii.gz), b-values (bval), b-vectors (bvec), an optional reversed phase encoded b=0 (rev_b0.nii.gz) and T1-weighted image (t1.nii.gz). The NIfTI format is chosen instead of DICOM: the naming convention is not the same across MRI sites in a DICOM. The user must extract the images, from the DICOM, in NIfTI format following the naming convention.

### 2.5. Diffusion-weighted imaging tasks

The DWI processing consists of 14 tasks (Figure 1.B) from raw DWI preprocessing steps, to the diffusion tensor imaging [Le Bihan et al., 2001] (DTI) metrics and fiber orientation distribution function [Descoteaux et al., 2007; Tournier et al., 2007] (fODF) metrics computation.

#### Denoising

This task is useful to remove the noise induced by the MRI acquisition, enhance the signal to noise ratio and improve the image quality and following metrics [Veraart et al., 2016]. The tool used is *dwidenoise* from MRtrix3. The denoising is performed in the original spatial resolution and uses the MP-PCA method [Veraart et al., 2016]. However, this denoising algorithm does not correct the Rician bias. This issue is a trade-off between computation time and denoising quality. In our experience, the majority of DWI datasets contain noise, which means that the denoising step is enabled by default. However, the denoising can be deactivated if the DWI is already denoised.

#### Eddy/Topup

First of all, we extract a brain mask in the *Bet Prelim DWI* task, with the *bet* [Smith, 2002] command. This first brain extraction permits to accelerate the Eddy/Topup computing only in the given brain mask. Topup corrects the brain deformation induced by the magnetic field susceptibility artefacts. Then, eddy corrects for the eddy-currents, corrects motion artefacts and applies a slice drop correction. In this pipeline, if a reversed phase encoded b=0 image is provided, the *topup* command is running on the b=0 and reversed phase encoded b=0 images to extract the deformation field. Then, the deformation field is applied and the *eddy* command is performed using the topup output. Eddy can be performed with two different commands: *eddy_openmp* and *eddy_cuda*. Like the denoising, Eddy/Topup is optional but enabled by default to enhance the quality of the DWI and correct the deformation bias. Moreover, the Eddy/-Topup processes are ran using only one thread to ensure the reproducibility. Commands such as *bet*, *topup* [Andersson et al., 2003; Smith et al., 2004] and *eddy* [Andersson and Sotiropoulos, 2016] come from the FSL package.

#### Brain Extraction

The brain mask is extracted with the *bet* [Smith, 2002] command from FSL. First, the extraction is performed on the b=0mm^2^/s image. Then the brain mask is applied to the whole DWI. The brain extraction is performed after Eddy/Topup to have a distortion-free brain mask corrected by Topup. This brain extraction is required to remove the skull and prepare the DWI to the T1 registration.

#### N4 bias correction

Typically, in brain acquisitions, the middle of the brain is less intense than its outer boundary. The N4 bias correction normalizes the image intensities and reduce this intensity bias. The N4 correction is computed after Eddy/Topup not to modify the intensity values that are important for the deformation bias and the eddy current correction. The N4 correction [Tustison et al., 2010] is performed on the b=0mm^2^/s with the ANTs command (*N4BiasFieldCorrection*). Then, the bias field is applied to the whole DWI [Shen et al., 2017; Raffelt et al., 2017; Mito et al., 2018; Constanzo et al., 2018].

#### Crop

After the N4 correction, the DWI is cropped to reduce the bounding box of the DWI. The function used here comes from DIPY package. This is simply to reduce the size of the image and accelerate following processing steps.

#### Normalize

The DWI is normalized to have a mean value in the WM of approximately 1000 [Raffelt et al., 2017; Mito et al., 2018]. This task permits analyzing datasets from different MRI scanners with the same acquisition scheme. The script is *dwinormalise* and is availabled in MRtrix3.

#### Resample

In the pipeline, the DWI is resampled to 1mm isotropic spatial resolution, which is usually the spatial resolution of the T1. The resample improves the anatomical details of the DWI [Dyrby et al., 2014] and has been used in several recent tractography algorithms [Girard et al., 2014; Smith et al., 2012]. This spatial resolution is modifiable in the configuration file. Moreover, the resample task is optional but enable by default. The functions used here are available in DIPY.

#### DTI metrics

DTI metrics are used in a large number of dMRI studies. The metrics computed from the diffusion tensor reconstruction are the axial diffusivity (AD), fractional anisotropy (FA), geodesic anisotropy (GA) [Batchelor et al., 2005], mean diffusivity (MD), radial diffusivity (RD), tensor, tensor norm [Kindlmann et al., 2007], tensor eigen-values, tensor eigenvectors, tensor mode, color FA [Pajevic and Pierpaoli, 1999]. DTI being a low b-value model, only the low b-values shells are extracted from the DWI for DTI fitting. We recommend to use b-values under b=1200mm^2^/s [Jensen and Helpern, 2010]. The b-values are extracted from the DWI (*Extract DTI shells* task) to compute the tensor model and extract the DTI metrics using DIPY [Garyfallidis et al., 2014a].

#### fODF metrics

Suitable shells are extracted from the *Extract fODF shells* task. We recommend to extract the b-values above b=700mm^2^/s. The fiber response function (FRF) used can be the average of all FRF across the subjects or set for all subjects. These options are set in the configuration file. The fODF, using constrained spherical deconvolution (CSD) [Descoteaux et al., 2007; Tournier et al., 2007], and the metrics are computed using fiber response function. The fODF metrics computed are the total and maximum Apparent Fiber Density (AFD) [Raffelt et al., 2012], the Number of Fiber Orientation (NuFO) [Dell’Acqua et al., 2013] and the principal fODFs orientations (up to 5 per voxel).

### 2.6. T1 weighted image tasks

The T1 tasks break down into 8 tasks (see Figure 1.C) from the raw T1 to the segmented tissues maps required for the tractography. The use of the *Denoising*, *N4 bias correction*, *Resample*, *Brain Extraction* and *Crop* tasks are the same as presented in the DWI tasks.

First, the T1 is denoised with the non-local means with rician correction method [Coupé et al., 2008]. The script implementation comes from the DIPY package. Then, the N4 bias correction [Tustison et al., 2010] (*N4BiasFieldCorrection*) from ANTs is performed. If necessary, the T1 is resampled at a given spatial resolution with DIPY. By default, the T1 is resampled to 1mm isotropic but this task can be disabled. After that, the brain mask is extracted with ANTs. To perform this brain extraction, the *antsBrainExtraction.sh* command is called. This brain extraction method is based on a T1 template that must be given in input of the pipeline. This template approach is more robust than the *bet* command. In our experience, *bet* command could remove a layer of gray matter that is more robust with the ANTs brain extraction. However, *antsBrainExtraction.sh* command has a longer runtime. Then, to reduce the bounding box of the image, the T1 is cropped with DIPY.

The three T1-specific tasks are the registration on the DWI, the tissue segmentation and the computation of the tracking masks.

#### Registration

After T1 preprocessing, the T1 image is registered [Avants et al., 2008] on the b=0mm^2^/s and the fractional anisotropy (FA) images. The registration is performed with nonlinear SyN ANTs (*antsRegistration*) multivariate option, where there are two (2) target images (b0 and FA) and the moving T1 image. This step is required to have the partial volume maps of the tissues used for the tracking task.

#### Tissue segmentation and tracking maps

Next, the white matter (WM), gray matter (GM), and cerebrospinal fluid partial volume maps are extracted with *fast* [Zhang et al., 2001] from the FSL package. These maps are used to compute the tracking maps [Girard et al., 2014]: inclusion and exclusion maps as well as seeding mask. Inclusion and exclusion maps are anatomical constraints for the anatomically-constrained tracking algorithm [Girard et al., 2014; Smith et al., 2012]. The seeding mask is the WM-GM interface or the WM mask.

### 2.7. Tracking task

The tracking process generates a set of streamlines that represent the underlying WM pathways. This object is called a tractogram [Côté et al., 2012, 2013]. The tractography algorithm used in this pipeline is the anatomically-constrained particle filter tracking [Girard et al., 2014]. This tracking uses the fODF image, an inclusion and an exclusion map and a seeding mask to generate the tractogram. This task compute a tractogram using the probabilistic or deterministic method. Here, we have two types of seeding: the total number of seeds or the number of seeds per voxel. By default, the probabilistic tractography is used with ten (10) seeds per voxel. Other parameters can be modified, such as the step size, maximum angle, and usual streamline-based tracking parameters, in the pipeline configuration file. By default the tracking process is reproducible using a fixed random seed number for the random number generator.

### 2.8. “Standard” pipelines

Most often, the reproducibility of dMRI pipeline results is never studied. However, this reproducibility aspect is important to have the same results over time. To obtain a baseline, a “standard” pipeline, based on the Mrtrix3 beginner tutorial, was ran 3 times.. The MRtrix3 pipeline is described in the Supplementary materials. Moreover, to perform other comparisons to a standard pipeline, our *TractoFlow* was ran 3 times without fixing the seed for the random number generator and the multithreading activated for *Eddy/Topup* task (called *TractoFlow-no-seeds*).

### 2.9. Reproducibility analysis

For each dataset, and each computer system, the pipeline was ran three times with the same parameters and the same Singularity container (commonly called test-test analysis). The parameters used across the pipeline are the default ones. Moreover, reproducibility of “standard” pipelines was computed on five subjects from *Penthera3T*.

#### Reproducibility measures of local diffusion and T1 maps

As shown in Figure 2, the correlation coefficient (CC) and mean absolute error (MAE) are assessed between 2 runs, between every diffusion metrics produced such as FA, diffusivities, total AFD. This amounts to a total of 3 values of CC and MAE per diffusion measure. Then, these 3 values are averaged to obtain a mean CC and mean MAE per subject, a total of 30 mean CCs and 30 mean MAEs. Finally, we average the 30 mean CCs and 30 mean MAEs to obtain the “global” mean CC and mean MAE across the datasets.

**Figure 2:**
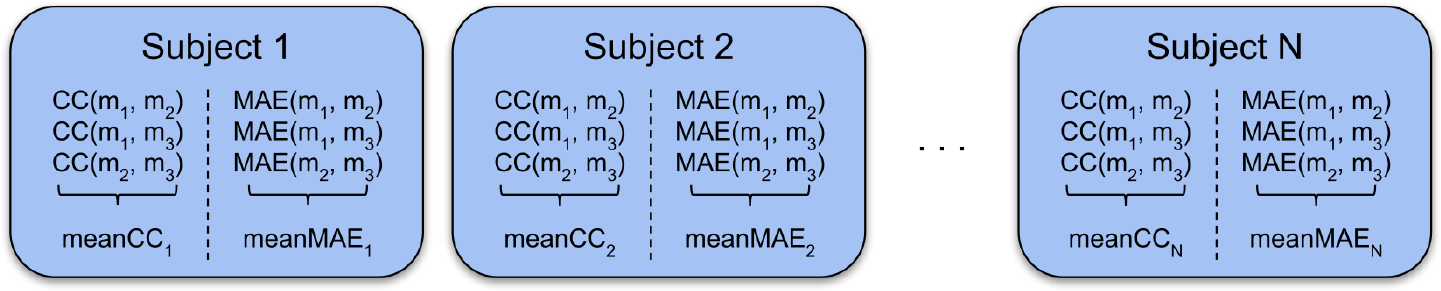
For each subject, the correlation coefficient (CC) and the mean absolute error (MAE) are computed for each map m*i* between runs *i*. The mean CC and mean MAE is obtained from 3 comparisons.

#### Tractogram reproducibility metrics

To evaluate the global reproducibility of the tractograms, the percentage of same segments of streamlines is computed between 2 runs. To obtain the reproducibility of the tractograms across the 30 subjects, the percentage of same segments per streamline is averaged. Moreover, a density map, that reports the number of streamlines per voxel, and an endpoints map, that reports the number of streamline endpoints per cortical voxel, are computed from the whole brain tractogram. Figure 3 shows the variability impact of the tissue maps on the tractogram, density map and endpoint map. The endpoints map evaluates only the endpoints (2 points only) of each streamline. Hence, the reproducibility of the endpoints map is expected to be more variable than the density map. These two maps are important to evaluate the reproducibility of the core of the streamlines (density map) and the reproducibility of the connectivity (endpoint maps) signature of streamlines. Finally, to evaluate the reproducibility of the density and endpoints maps, a mean CC and mean MAE is computed on each subject across the 3 runs. Finally, the mean CCs and mean MAEs of each subject is averaged.

**Figure 3:**
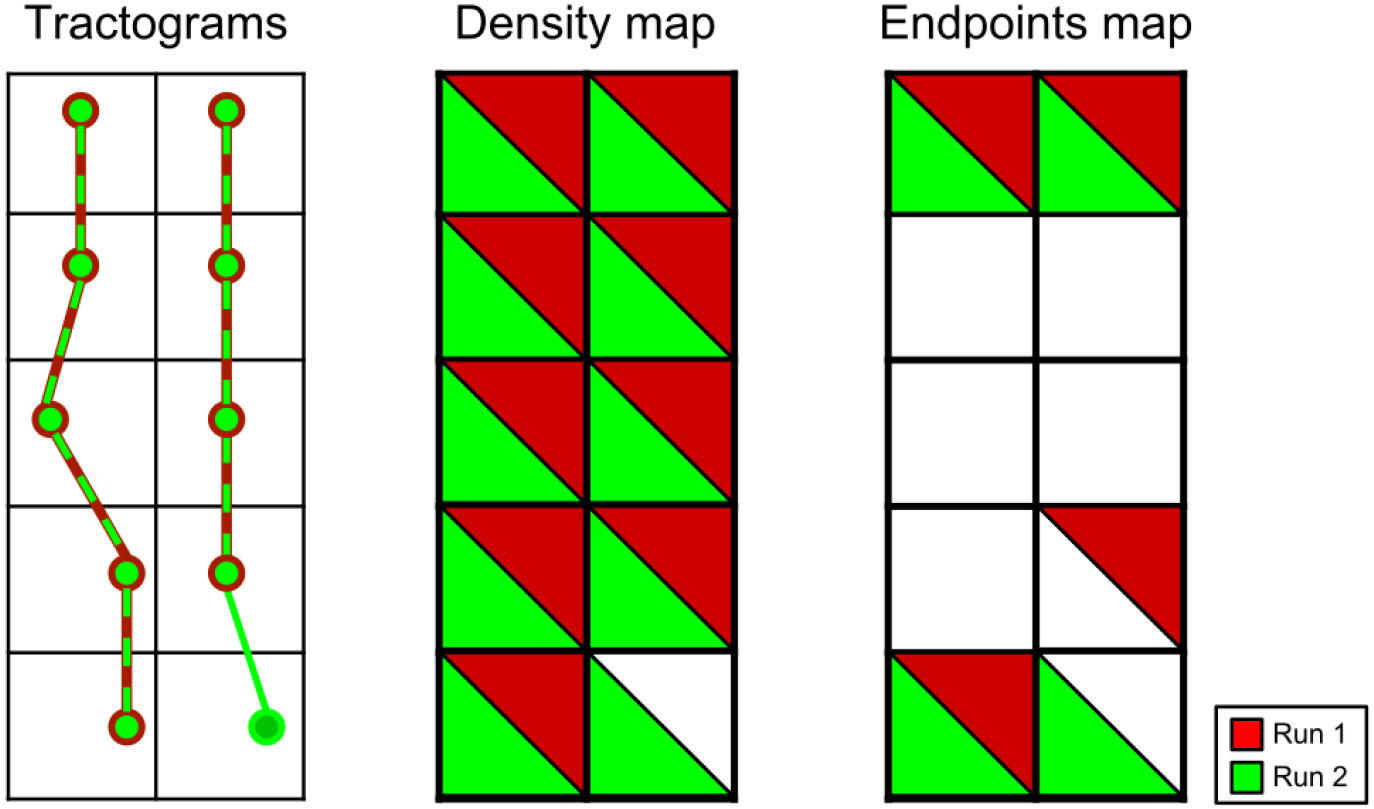
For two runs, in red the first run and in green the second run, we represent the tractograms (left), the density map (center) and the endpoints map (right).

Then, a connectivity matrix is computed for each subject and each run (see Figure 4.A). A connectivity matrix is based on a atlas that contains parcels. The matrix reports the number of streamlines that connect from a parcel to another for each parcel combinations. This is the usual structural connectivity matrix used in connectomics [Hagmann et al., 2008]. The connectivity matrix computed for the tractogram reproducibility analysis is based on the Brainnetome atlas [Fan et al., 2016]. The Brainnetome atlas must be registered to each subject. To have a reproducible registration, we create a template between the three runs for each subject and the registration is performed on the template. This technique ensures using the same atlas for three runs. Thus, we only evaluate the reproducibility of the pipeline, without introducing biases and errors from the atlas registration. The mean CC and the mean MAE across the 3 runs are used to evaluate the reproducibility of the connectivity matrix. Then, the 30 mean CCs and mean MAEs are respectively averaged to obtain a “global” average across the datasets.

**Figure 4:**
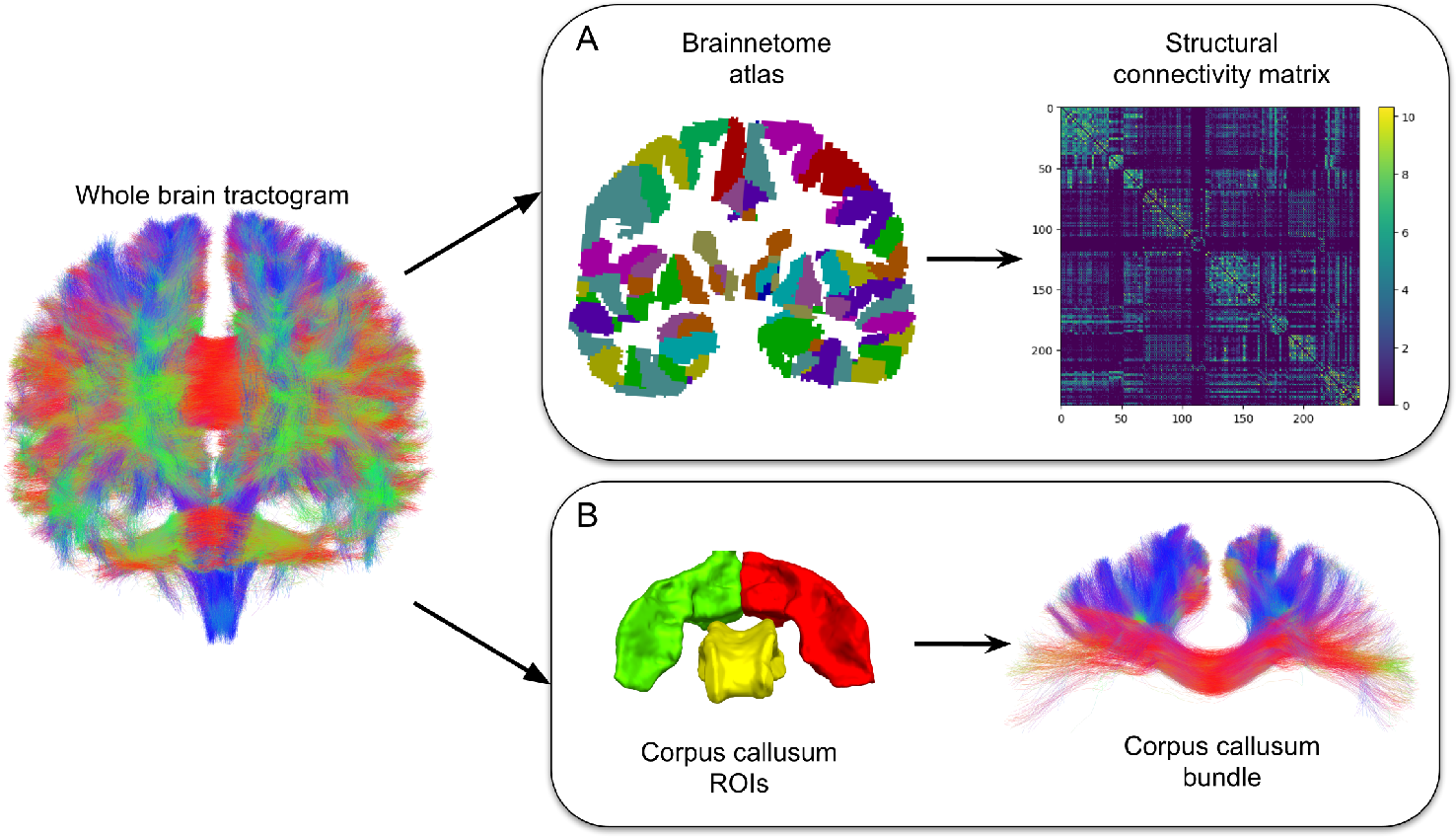
For each subject and each run, the structural connectivity matrix is obtained from the Brainnetome atlas and the whole brain tractogram (A). Corpus callusum bundle is obtained from region of interests and the whole brain tractogram (B).

Finally, to quantify the reproducibility at the WM bundle level, the mid-body of the corpus callosum bundle, that must connect the precentral and postcentral gyri, is extracted from the whole tractogram for each subject and each run using a common a region of interest (ROI) approach [Rheault et al., 2018] (see Figure 4.B). The ROIs are registered on each subject. As for the connectivity matrix atlas registration, a template across the three runs is computed to perform the same ROI registration for each subject. The percentage of same streamline segments is also computed as previously presented. Moreover, as explained for the whole tractogram, a density map and an endpoints map is computed for the corpus callusum and the reproducibility of these maps is evaluated.

## 3. Results

Measures presented in this section come from the *Cluster 1*. Similar trends and conclusions are obtained from other computers. Hence, we provide the full results from *Cluster 2* and the *Desktop* computers in the Supplementary materials.

### 3.1. Reproducibility of “standard” pipelines

For the “standard” pipelines, to evaluate the reproducibility for the whole brain and corpus callusum, the percentage of same streamline segments, density and endpoints map were computed on five subjects of the *Penthera3T*. Both the *TractoFlow-no-seeds* (without fixing seeds and allowing multi-threading) and the MRtrix pipeline presented in Subsections 2.8, are unreproducible. In terms of tractogram, none of the streamlines segments are similar between two runs for the exact same acquisition. This unreproducibility impacts the reproducibility of the maps extracted from the tractogram (see Table 2). The streamline density map has a mean CC and MAE of 0.99 and 0.04 respectively on the whole tractogram and, for the corpus callusum, the mean CC and mean MAE is 0.99 and 0.30 respectively. Then, the streamline endpoints map have a mean CC and MAE of 0.92 and 0.19 respectively on the whole tractogram and, for the corpus callosum, the mean CC and MAE dropped to 0.73 and 0.61 respectively. Finally, the connectivity matrix have a mean CC and MAE of 0.99 and 0.30 respectively. These results highlight the importance of fixing the seeds and the number of threads to use, to ensure reproducibility higher than 95%. These standard pipelines will be further analyzed and discussed later in the Discussion section.

**Table 2:** For the “standard” pipelines, the mean correlation coefficient (CC) and mean MAE are computed for the whole tractogram density and endpoints map, corpus callusum density and endpoints map and connectivity matrix.

|  |  | Mean CC (SD) | Mean MAE (SD) |
| --- | --- | --- | --- |
| Whole tractogram | Density map | 0.99711 (0.00024) | 0.04117 (0.00694) |
|  | Enpoints map | 0.92912 (0.01078) | 0.18733 (0.01770) |
| Corpus callosum | Density map | 0.99237 (0.00223) | 0.30456 (0.02222) |
|  | Endpoints map | 0.73004 (0.06718) | 0.60619 (0.03568) |
| Connectivity matrix |  | 0.99905 (0.00006) | 0.30184 (0.00756) |

Now to evaluate the reproducibility of our *TractoFlow*, the reproducibility of local diffusion and T1 maps are evaluated. Then, the reproducibility of the tractogram, corpus callusum and connectivity matrix is computed. Finally, the runtime of the pipeline is logged.

### 3.2. Reproducibility of local diffusion and T1 maps

First, the reproducibility of the different measures produced in the pipeline is analyzed. All the DWI tasks (*Preliminary brain extraction, Denoise, Eddy/Topup, Brain Extraction, N4, Crop, Normalize, Resample, Extract DTI and fODF shells, DTI metrics Compute FRF, Mean FRF, fODF metrics*) are 100% reproducible (i.e. mean CC of 1 and mean MAE of 0), hence not shown here. Only two T1 tasks introduce a small MAE with a high correlation coefficient, caused by the brain extraction and the registration steps, as seen in Table 3. Across the two datasets, the smallest mean CC is 0.99988 and the biggest mean MAE is 0.00063.

**Table 3:** Mean correlation coefficient (CC) and mean MAE for the brain extraction and registration task on the T1.

|  |  | Mean CC (SD) | Mean MAE (SD) |
| --- | --- | --- | --- |
| Penthera1.5T | Bet T1 | 0.99988 (0.00020) | 0.00006 (0.00011) |
|  | Register T1 | 0.99995 (0.00008) | 0.00051 (0.00076) |
| Penthera3T | Bet T1 | 0.99993 (0.00008) | 0.00016 (0.00019) |
|  | Register T1 | 0.99997 (0.00003) | 0.00063 (0.00057) |

As expected, errors generated from the T1 brain extraction and registration have an impact on the *Tissue segmentation* task. Table 4 highlights the mean CC and mean MAE for WM, GM and CSF maps. These small differences are due to the probabilistic classification of the tissues. This probability is slightly different in the partial volume and introduces these errors due to a numerical instability of the *Registration* task. However, the mean CC is 0.99 or higher and the mean MAE 0.015 or lower.

**Table 4:** Mean correlation coefficient (CC) and mean MAE for WM, GM and CSF maps extracted from the T1 registered.

|  |  | Mean CC (SD) | Mean MAE (SD) |
| --- | --- | --- | --- |
| Penthera1.5T | WM map | 0.99734 (0.00211) | 0.00168 (0.00152) |
|  | GM map | 0.99701 (0.00264) | 0.00186 (0.00164) |
|  | CSF map | 0.98657 (0.01190) | 0.00442 (0.00401) |
| Penthera3T | WM map | 0.99734 (0.00211) | 0.00168 (0.00152) |
|  | GM map | 0.99701 (0.00264) | 0.00186 (0.00164) |
|  | CSF map | 0.98657 (0.01190) | 0.00442 (0.00401) |

### 3.3. Tracking reproducibility

*Whole tractogram*. The percentage of same segments of streamlines for Penthera1.5T and Penthera3T were 98.5% and 97.7% respectively. Table 5 shows the mean correlation coefficient and the mean MAE for the density map and endpoint map.

**Table 5:** Average on all subjects and runs of the Mean Absolute Error (MAE) and correlation coefficient (CC) on the density map and endpoint maps of the whole tractogram.

|  |  | Mean CC (SD) | Mean MAE (SD) |
| --- | --- | --- | --- |
| Penthera1.5T | Density map | 0.99996 (0.00006) | 0.00271 (0.00371) |
|  | Endpoint map | 0.99767 (0.00325) | 0.01111 (0.01503) |
| Penthera3T | Density map | 0.99991 (0.00013) | 0.00423 (0.00369) |
|  | Endpoint map | 0.99668 (0.00340) | 0.01687 (0.01411) |

Across Penthera1.5T and Penthera3T, the lowest mean CC for the density map is 0.99991 while for the endpoint map is 0.99668. For the mean MAE, the same pattern is observed. These small errors come from the small differences in the WM, GM and CSF mask introduced by the brain extraction and registration process, which impact the anatomically-constrained particle filter tracking algorithm [Girard et al., 2014] that forces streamlines to stop in the GM map.

#### Connectivity matrix

As shown in Table 6, the reproducibility of the connectivity matrix is high with a mean CC of 0.99999 across the two datasets. Moreover, the mean MAE is low with values of 0.01125 and 0.01830 for Penthera1.5T and Penthera3T respectively.

**Table 6:**
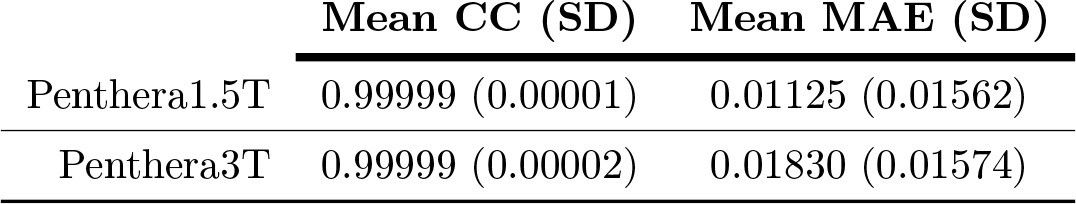
Average on all subjects and runs of the Mean Absolute Error (MAE) and correlation coefficient (CC) on the connectivity matrix.

|  | Mean CC (SD) | Mean MAE (SD) |
| --- | --- | --- |
| Penthera1.5T | 0.99999 (0.00001) | 0.01125 (0.01562) |
| Penthera3T | 0.99999 (0.00002) | 0.01830 (0.01574) |

#### Corpus callosum bundle

The percentage of same segments of streamlines for the corpus callusum on the Penthera1.5T and Penthera3T were 94.1% and 93.3% reproducible respectively. Figure 5.C shows the streamline segments that are different between two runs. Corpus callusum volume has a difference of 0.07% and a difference of 0.39% for the number of streamlines.

**Figure 5:**
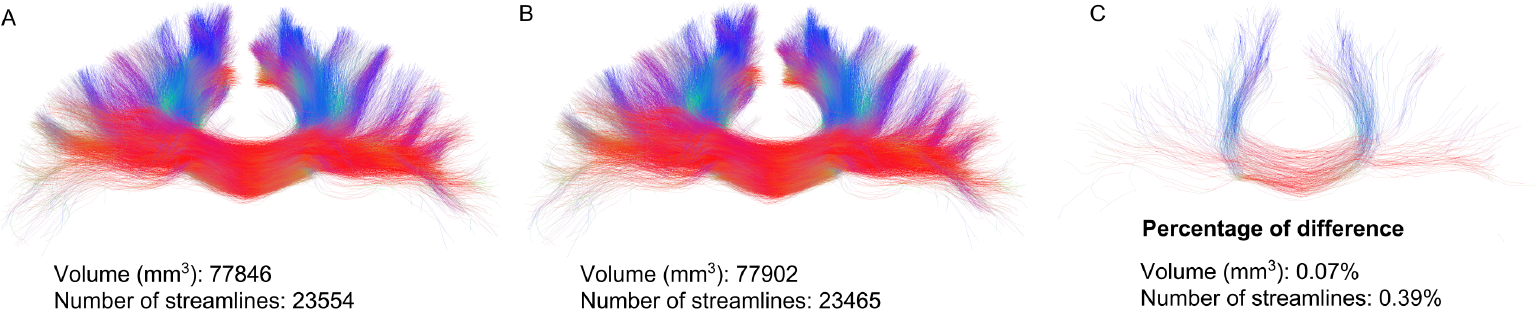
In A, the corpus callusum extracted from a subject in the run 1. In B, the corpus callusum extracted from the same subject in the run 2. In C, the fibers not reproducible in the corpus callusum.

Second, Table 7 summarizes the mean CC and mean MAE across the 30 subjects for two datasets. The density map mean CC is 0.99757 and 0.99868 respectively for Penthera1.5T and Penthera3T. The mean CC for the endpoints map are significantly lower than the values observed for the whole tractogram. For Penthera1.5T and Penthera3T, the mean CC for the endpoints map is respectively 0.95666 and 0.91487.

**Table 7:** Average on all subjects and runs of the Mean Absolute Error (MAE) and correlation coefficient (CC) on the density map and endpoint maps of the corpus callusum.

|  |  | Mean CC (SD) | Mean MAE (SD) |
| --- | --- | --- | --- |
| Penthera1.5T | Density map | 0.99757 (0.00657) | 0.02445 (0.03581) |
|  | Endpoint map | 0.95666 (0.06219) | 0.07897 (0.11094) |
| Penthera3T | Density map | 0.99868 (0.00257) | 0.02708 (0.02924) |
|  | Endpoint map | 0.91487 (0.10748) | 0.10231 (0.10608) |

The mean CC values observed for the corpus callusum are lower than the values for the whole tractogram for the density and endpoints map. These differences have the same explanation as the whole tractogram (see Figure 3). However, the corpus callusum have less streamlines than a whole tractogram and the streamlines segment different across the runs have a bigger impact.

### 3.4. Benchmark

An important part of the pipeline is the fast runtime. For each run of 30 subjects, the runtime was logged. Table 8 reports the runtime for each run for two datasets. The average runtime for Penthera1.5T and Penthera3T were 17 hours 12 minutes and 16 hours and 37 minutes respectively.

**Table 8:** Time of computing for each run and each dataset (30 subjects).

|  | Run 1 | Run 2 | Run 3 | Mean (SD) |
| --- | --- | --- | --- | --- |
| Penthera1.5T | 17h07 | 17h35 | 16h54 | 17h12 (0.015h) |
| Penthera3T | 16h33 | 16h22 | 16h56 | 16h37 (0.012h) |

For the HCP subset, the pipeline was ran one time without the *Eddy/Topup*, *resample T1/DWI* tasks. The seeding mask used was the white matter mask. The tracking algorithm is the probabilistic method with 30 seeds per voxel. The pipeline was launched on 4 cluster nodes of 32 threads and 125 Gb of memory for each node. The runtime for the 105 subjects was approximately 25 hours.

## 4 Discussion

Our *TractoFlow* allows processing dMRI datasets efficiently while ensuring the reproducibility of the local DTI/HARDI measures, T1 processing steps and tractography results with little to no installation steps. A goal of this pipeline is to promote efficient and reproducible diffusion tractography processing for open science, with industry standards. *TractoFlow* is open and available (https://github.com/scilus/tractoflow) as well as Penthera1.5T (https://doi.org/10.5281/zenodo.2602022) and Penthera3T (https://doi.org/10.5281/zenodo.2602049).

### 4.1. Nextflow and Singularity

Nextflow was the pipeline tool chosen for the *TractFlow*. However, other pipelining tools exist such as Nipype [Gorgolewski et al., 2011] or Luigi (https://luigi.readthedocs.io). Nipype is a tool adapted for neuroscience processing. However, creating and maintaining Nipype pipelines is more complex than a Nextflow pipeline. Also, using and adding packages that are not available in Nipype is a process involving more steps. Luigi is not associated with neuroscience processing. However, as Nipype, it is harder to create a Luigi pipeline than a Nextflow pipeline. Finally, Snakemake [Köster and Rahmann, 2012] is a pipeline creation tool similar to Nextflow.

As Singularity, Docker (https://www.docker.com/) is another container software. However, Docker requires administrator permissions, which is not realistic for HPC deployment, while Singularity requires no such permissions. A Docker container is available for OSX users with the same packages as the Singularity container.

### 4.2. Pipeline processing steps

Using Nextflow, the pipeline is easy to modify and upgrade over time. The code is designed so that one task does one specific processing step. As an example, Figure 6 shows the code of the DWI denoising task in the *TractoFlow* Nextflow pipeline. As previously introduced, our denoising method does not support Rician bias. Our denoising can be easily replaced by another one supporting the Rician bias in the *script* scope (see Figure 6), such as non local mean and Non Local Spatial and Angular Matching (NLSAM) [Descoteaux et al., 2008; St-Jean et al., 2016]. These denoising techniques robust to rician noise and spatially-varying noise, such as NLSAM, typically require more assumptions about the data (number of coils used, non-masked background in the image, amongst others), which make them less scalable across subjects, scanners, sites. However, as these techniques evolve, become more robust and more efficient, they could easily be integrated in the TractoFlow pipeline.

**Figure 6:**
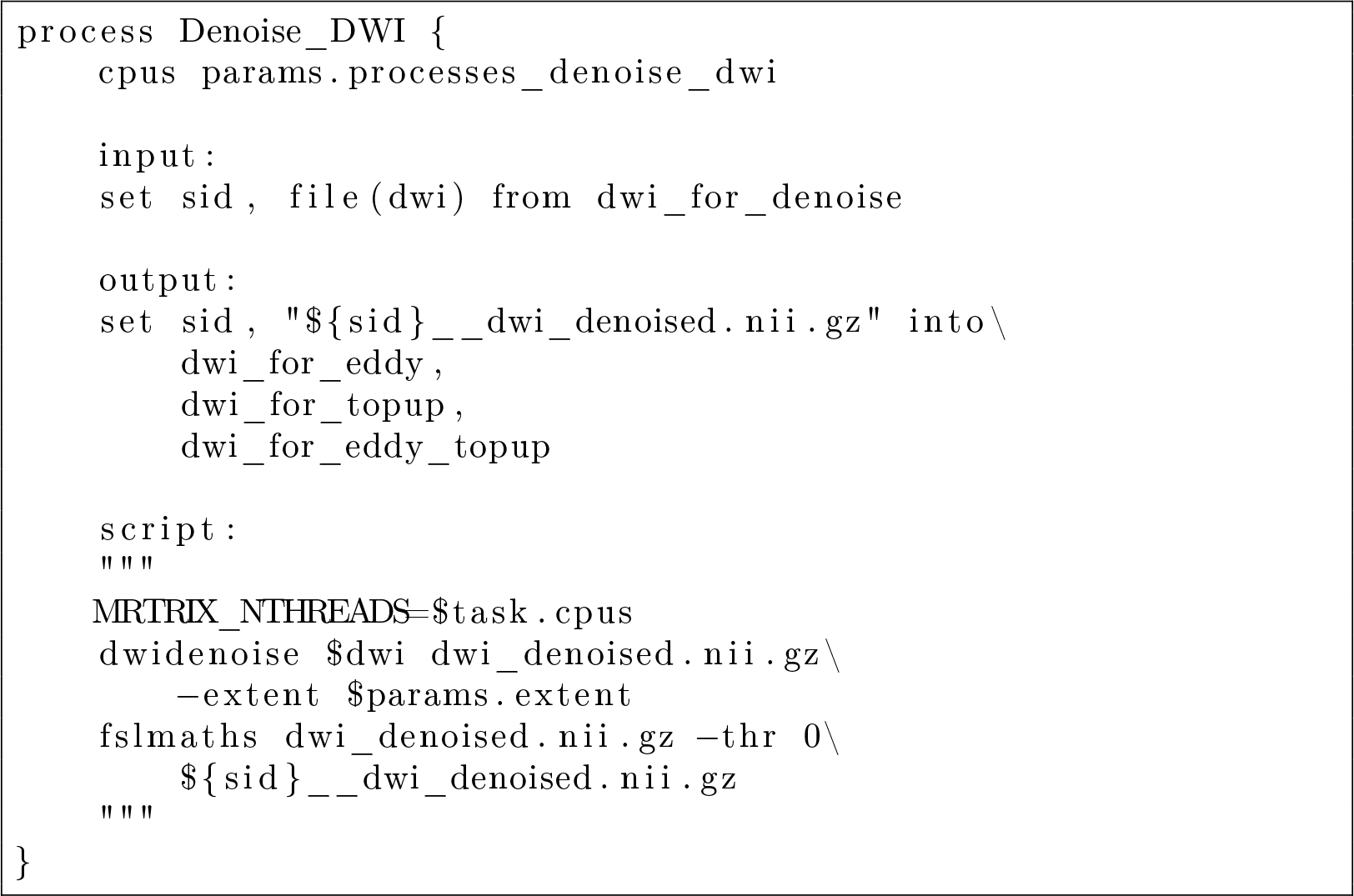
Example of a *TractoFlow* task: DWI denoising. In the *input* scope the files requiered by the task. In the *output* scope, the DWI denoised. In the *script* scope, the denoising command to run: *dwidenoise*.

Moreover, a Gibbs ringing correction task must be added to the pipeline in the future. The MRtrix command *mrdegibbs* [Kellner et al., 2016] must be tested on multiple datasets to check the reproducibility of the command and the efficiency in terms of results and runtime. On our tested datasets, Gibbs ringing was barely visible.

Moreover, currently, fODFs are computed using a classical CSD approach implemented in Dipy. This was chosen as it supports both multi-shell and single-shell acquisition as input [Garyfallidis et al., 2014b]. However, other cutting-edge and more optimal methods exist, such as multi-tissues multi-shells CSD [Jeurissen et al., 2014] or three tissues single shell [Dhollander and Connelly, 2016]. As for the DWI denoising, the command used can be easily modified to support a more cutting-edge fODF processing. Three tissue single-shell and multi-shell multi-tissue methods are not yet implemented in Dipy but are part of ongoing developments in the Dipy community.

Tractography is the last processing step of the pipeline. It was a deliberate choice to stop processing at fiber tracking. Separate Nextflow pipelines are currently under construction for fiber bundling, tractometry, statistics, and more. On the other hand, tractography could also become optional in the TractoFlow pipeline if, for example, users are only interested in DTI/HARDI measures.

### 4.3. Reproducibility analysis

Our *TractoFlow* pipeline is a first step to unify dMRI processing. The pipeline is almost 100% reproducible, with an average tractogram reconstruction reproducibility across the 3 runs of 98.1%. All reproducibility analysis performed in this study shows a minimum correlation coefficient of 0.9. As shown in the supplementary materials, the pipeline is also reproducible on the desktop and the cluster 2. Nonetheless, the pipeline can reach 100% reproducibility with a 50% slower runtime. The brain extraction and registration tasks must then be ran in single-thread mode. The runtime for the 30 subjects of Penthera1.5T takes 26 hours and 42 minutes instead of 17 hours. This modification ensures the 100% reproducibility of the maps generated across the pipeline, the tracking and the metrics and maps generated from the tracking.

In terms of “standard” pipelines, it is not surprising that both our *TractoFlow-no-seeds* and MRtrix are not reproducible in track segments. The seeds are random and not the same from run to run. Hence, initial points vary slightly, which will yield different track segments over the whole streamline. Table 9 shows the differences of mean CC for the “standard” pipelines and *TractoFlow*. In terms of tractometry, tract-based analysis and connectomics, this can have a sizeable impact, as we have showed that endpoints maps, connectomes and bundles are not reproducible. This can lead to unreproducible results in experiments and in time, which can affect conclusions and interpretations of results in application studies.

**Table 9:** Mean correlation coefficient comparison between “standard” pipelines and *TractoFlow* (Penthera3T database), for the whole tractogram density and endpoints map, corpus callusum density and endpoints map and connectivity matrix.

|  |  | "Standard" pipelines | *TractoFlow* |
| --- | --- | --- | --- |
| Whole tractogram | Density map | 0.99711 | 0.99991 |
|  | Endpoints map | 0.92912 | 0.99668 |
| Corpus callosum | Density map | 0.99237 | 0.99868 |
|  | Endpoints map | 0.73004 | 0.91487 |
| Connectivity matrix |  | 0.99905 | 0.99999 |

### 4.4. Advantages of a High Performance Computer

A High Performance Computer (HPC) and our Nextflow pipeline enables processing large number of subjects at the same time with only a single command. Using a Singularity container, tedious setup of libraries and processing environment becomes effortless. The container image is simply given as input to the Nextflow pipeline. Moreover, if the HPC supports it, the pipeline can run on multiple nodes, the number of subjects ran at the same time is larger and the runtime is reduced. This feature depends on the setup of the HPC, but is usually enabled effortlessly in Nextflow. Moreover, the benchmarking shows the reproducibility of time. This runtime reproducibility is important in HPC context when a walltime, a maximum runtime value, must be set before launching the pipeline.

In our experience and in our own lab, we recommend the 100% reproducible version for a HPC use, as Nextflow is very good at optimizing job dispatching to maximize threads used in the processing. Since the number of threads is typically much higher than on a personal desktop, this does not affect run time too badly. On the other hand, if the pipeline is running on a personal desktop, we recommend the default parameters, which lead to 98% reproducibility in a faster time.

### 4.5. Future work

Currently, supporting the Brain Imaging Data Structure (BIDS) [Gorgolewski et al., 2016] as an input structure option is almost done. Moreover, to promote open-science, *TractoFlow* will be available on CBRAIN [Sherif et al., 2014]. CBRAIN is a cloud service to process data, similar to Brainlife. In Brainlife, some pipeline exist to process tractography [Avesani et al., 2018].

Some large datasets such as the UK Biobank [Sudlow et al., 2015] dataset will be processed with our TractoFlow pipeline. The pipeline will run on Compute Canada HPC on multiple nodes. Moreover, it is part of future work to replace some processing steps of the pipeline using machine learning algorithms. The T1 brain extraction and the registration [Balakrishnan et al., 2018] steps can be accelerated, improved and made more reproducible with a machine learning algorithms.

Moreover, a quality check (QC) and quality assurance (QA) pipeline is in development to evaluate the results computed from this pipeline. This QC/QA pipeline will be a Nextflow pipeline and will generate HTML reports for each step of the processing pipeline presented in this paper. The reports contain screenshots, statistics and basic outlier detection routine implemented at every critical step of the pipeline.

## 5. Conclusion

We proposed a new Nextflow pipeline to robustly and efficiently process dMRI datasets (big or small) with little to no setup, thanks to the use of a Singularity container. The container and Nextflow parameterization of all steps ensure the reproducibility at 98% of the results over time. Moreover, the pipeline is fast: i) 17 hours for 30 subjects with a standard 2mm isotropic DWI with 64 directions and ii) 25 hours for 105 subjects from HCP dataset. If needed, the pipeline can have a 100% reproducibility score with a 50% longer runtime. If a high performance computer is available, we highly recommend running the pipeline with the 100% reproducible configuration. The un-reproducibility of a “standard” pipeline can impact analysis, bias results over time and, as a result, make dMRI quantitative analysis un-usable for clinical trials. The reproducibility issue of dMRI pipelines must be taken into account by the community at the era of open science and large open databases. Our new TractoFlow dMRI tractography pipeline is a first step towards robust, fast and reproducible white matter analysis, which is available here (https://github.com/scilus/tractoflow).

## 6. Acknowledgements

Thank you to Jean-Christophe Houde, Arnaud Boré, Francois Rheault and Felix Morency for their insight and help on this work. Thank you to Jon Haitz Legarreta for his proofreading. A special thank you to MITACS for funding this work through the Mitacs Accelerate program. Thank you to the Neuroinformatics Chair of the Sherbrooke University which helped push forward neurosciences research.

