## Supplementary materials for "TractoFlow: A robust, efficient and reproducible diffusion MRI pipeline leveraging Nextflow & Singularity"

### 1. Supplementary materials

#### *1.1. Tools version*

- FSL: version 5.0.11
- Mrtrix3: version 3.0_RC3
- ANTs: version 2.3.1
- DIPY: version 0.16

#### *1.2. Configuration file*

```
//**Global options**//
b0_thr_extract_b0=10
dwi_shell_tolerance=20

//**Preliminary DWI brain extraction**//
dilate_b0_mask_prelim_brain_extraction=5
bet_prelim_f=0.16

//**Denoise dwi (dwidenoise in Mrtrix3)**//
run_dwi_denoising=true
extent=7

//**Topup**//
run_topup=true
config_topup="b02b0.cnf"
encoding_direction="y"
dwell_time=0.062
prefix_topup="topup_results"

//**Eddy**//
run_eddy=true
eddy_cmd="eddy_openmp"
bet_topup_before_eddy_f=0.16
use_slice_drop_correction=true

//**Final DWI BET**//
bet_dwi_final_f=0.16

//**Resample T1**//
run_resample_t1=true
t1_resolution=1
t1_interpolation="lin"

//**Resample DWI**//
run_resample_dwi=true
```

```
dwi_resolution=1
dwi_interpolation="cubic"

//**DTI shells**//
dti_shells="0 300 1000"

//**fODF shells**//
fodf_shells="0 1000 2000"

//**Segment tissues**//
number_of_tissues=3

//**Compute fiber response function (frf)**//
fa=0.7
min_fa=0.5
min_nvox=300
roi_radius=20
set_frf=false
manual_frf="15,4,4"

//**Mean fiber response function (frf)**//
mean_frf=true

//**Compute fODF metrics**//
sh_order=8
basis="descoteaux07"
fodf_metrics_a_factor=2.0
relative_threshold=0.1
max_fa_in_ventricle=0.1
min_md_in_ventricle=0.003

//**Seeding mask**//
wm_seeding=true

//**PFT tracking**//
compress_streamlines=true
algo="prob"
seeding="npv"
nbr_seeds=10
random=0
step=0.5
theta=20
sfthres=0.1
sfthres_init=0.5
min_len=10
max_len=300
particles=15
```

```
back=2
front=1
compress_value=0.2

//**Number of processes per tasks**//
processes_brain_extraction_t1=8
processes_denoise_dwi=4
processes_denoise_t1=4
processes_eddy=1
processes_fodf=4
processes_registration=4

//**Template T1 path**//
template_t1="/human-data/mni_152_sym_09c/t1"
```

*1.3. Mrtrix3 pipeline*

```
#!/bin/bash

subj_id=../Subj_0
dwi=${subj_id}/dwi.nii.gz
bval=${subj_id}/bval
bvec=${subj_id}/bvec

dwidenoise $dwi dwi_denoised.nii.gz -force

dwi=dwi_denoised.nii.gz

dwipreproc -rpe_none -cuda -pe_dir AP $dwi dwi_preproc.nii.gz \
-fslgrad $bvec $bval -force

dwi=dwi_preproc.nii.gz

dwi2mask $dwi dwi_mask.nii.gz -fslgrad $bvec $bval -force

dwi2response tournier $dwi frf.txt -mask dwi_mask.nii.gz \
-fslgrad $bvec $bval -force

dwi2fod csd $dwi frf.txt fodf.nii.gz -mask dwi_mask.nii.gz \
-fslgrad $bvec $bval -force

tckgen fodf.nii.gz tracking.tck -seed_image dwi_mask.nii.gz \
-mask dwi_mask.nii.gz -select 1000000 -force
```

*1.4. Reproducibility results: Cluster 2*

*1.4.1. Maps reproducibility*

| | | **Mean CC (SD)** | **Mean MAE (SD)** |
|---|---|---|---|
| Penthera1.5T | Bet T1 | 0.99984 (0.00024) | 0.00009 (0.00013) |
| | Register T1 | 0.99994 (0.00009) | 0.00061 (0.00074) |
| | WM map | 0.99763 (0.00275) | 0.00134 (0.00159) |
| | GM map | 0.99753 (0.00291) | 0.00144 (0.00170) |
| | CSF map | 0.98770 (0.01464) | 0.00410 (0.00484) |
| Penthera3T | Bet T1 | 0.99991 (0.00010) | 0.00018 (0.00021) |
| | Register T1 | 0.99997 (0.00003) | 0.00054 (0.00055) |
| | WM map | 0.99783 (0.00198) | 0.00137 (0.00134) |
| | GM map | 0.99753 (0.00233) | 0.00154 (0.00146) |
| | CSF map | 0.98858 (0.01134) | 0.00368 (0.00360) |

Table 1: Mean correlation coefficient (CC) and mean MAE for the brain extraction task and registration task on the T1.

*1.4.2. Whole tractogram reproducibility*

| | | **Mean CC (SD)** | **Mean MAE (SD)** |
|---|---|---|---|
| Penthera1.5T | Density map | 0.99995 (0.00007) | 0.00327 (0.00373) |
| | Endpoint map | 0.99720 (0.00345) | 0.01330 (0.01476) |
| Penthera3T | Density map | 0.99993 (0.00009) | 0.00357 (0.00325) |
| | Endpoint map | 0.99692 (0.00391) | 0.01436 (0.01257) |

Table 2: Average on all subjects and runs of the Mean Absolute Error (MAE) and correlation coefficient (CC) on the density map and endpoint maps of the whole tractogram.

*1.4.3. Bundle reproducibility*

| | | **Mean CC (SD)** | **Mean MAE (SD)** |
|---|---|---|---|
| Penthera1.5T | Density map | 0.99803 (0.00569) | 0.02842 (0.03631) |
| | Endpoint map | 0.94913 (0.06665) | 0.09116 (0.11090) |
| Penthera3T | Density map | 0.99826 (0.00447) | 0.02481 (0.02938) |
| | Endpoint map | 0.91934 (0.11554) | 0.09417 (0.10567) |

Table 3: Average on all subjects and runs of the Mean Absolute Error (MAE) and correlation coefficient (CC) on the density map and endpoint maps of the corpus callusum.

##### *1.4.4. Connectivity matrix reproducibility*

| | Mean CC (SD) | Mean MAE (SD) |
|---|---|---|
| Penthera1.5T | 0.99999 (0.00001) | 0.01363 (0.01552) |
| Penthera3T | 0.99999 (0.00001) | 0.01629 (0.01491) |

Table 4: Average on all subjects and runs of the Mean Absolute Error (MAE) and correlation coefficient (CC) on the connectivity matrix.

#### *1.5. Reproducibility results: Desktop*

##### *1.5.1. Maps reproducibility*

| | | Mean CC (SD) | Mean MAE (SD) |
|---|---|---|---|
| Penthera1.5T | Bet T1 | 0.99987 (0.00020) | 0.00007 (0.00011) |
| | Register T1 | 0.99994 (0.00009) | 0.00068 (0.00095) |
| | WM map | 0.99755 (0.00319) | 0.00139 (0.00179) |
| | GM map | 0.99737 (0.00355) | 0.00160 (0.00220) |
| | CSF map | 0.98656 (0.01844) | 0.00446 (0.00614) |
| Penthera3T | Bet T1 | 0.99991 (0.00011) | 0.00018 (0.00023) |
| | Register T1 | 0.99997 (0.00004) | 0.00061 (0.00058) |
| | WM map | 0.99746 (0.00227) | 0.00157 (0.00134) |
| | GM map | 0.99720 (0.00241) | 0.00174 (0.00148) |
| | CSF map | 0.98719 (0.01175) | 0.00420 (0.00367) |

Table 5: Mean correlation coefficient (CC) and mean MAE for the brain extraction task and registration task on the T1.

##### *1.5.2. Whole tractogram reproducibility*

| | | Mean CC (SD) | Mean MAE (SD) |
|---|---|---|---|
| Penthera1.5T | Density map | 0.99995 (0.00008) | 0.00343 (0.00433) |
| | Endpoint map | 0.99720 (0.00389) | 0.01394 (0.01769) |
| Penthera3T | Density map | 0.99993 (0.00011) | 0.00396 (0.00306) |
| | Endpoint map | 0.99699 (0.00335) | 0.01588 (0.01193) |

Table 6: Average on all subjects and runs of the Mean Absolute Error (MAE) and correlation coefficient (CC) on the density map and endpoint maps of the whole tractogram.

#### *1.5.3. Bundle reproducibility*

| | | Mean CC (SD) | Mean MAE (SD) |
|---|---|---|---|
| Penthera1.5T | Density map | 0.99777 (0.00568) | 0.02804 (0.03901) |
| | Endpoint map | 0.94879 (0.06894) | 0.08756 (0.11744) |
| Penthera3T | Density map | 0.99827 (0.00471) | 0.02749 (0.03257) |
| | Endpoint map | 0.91600 (0.11292) | 0.09804 (0.10809) |

Table 7: Average on all subjects and runs of the Mean Absolute Error (MAE) and correlation coefficient (CC) on the density map and endpoint maps of the corpus callusum.

#### *1.5.4. Connectivity matrix reproducibility*

| | Mean CC (SD) | Mean MAE (SD) |
|---|---|---|
| Penthera1.5T | 0.99999 (0.00001) | 0.01343 (0.01787) |
| Penthera3T | 0.99999 (0.00001) | 0.01707 (0.01365) |

Table 8: Average on all subjects and runs of the Mean Absolute Error (MAE) and correlation coefficient (CC) on the connectivity matrix.

#### *1.6. Pipeline runtime for the cluster 2 and the desktop*

**Cluster 2**

| | Run 1 | Run 2 | Run 3 | Mean (SD) |
|---|---|---|---|---|
| Penthera1.5T | 29h14 | 29h48 | 24h03 | 26h42 (0.108h) |
| Penthera3T | 23h03 | 26h17 | 26h15 | 25h11 (0.077h) |

Table 9: Time of computing for each run and each dataset.

**Desktop**

| | Run 1 | Run 2 | Run 3 | Mean (SD) |
|---|---|---|---|---|
| Penthera1.5T | 114h45 | 118h13 | 115h20 | 116h06 (0.077h) |
| Penthera3T | 116h28 | 114h04 | 114h27 | 115h00 (0.054h) |

Table 10: Time of computing for each run and each dataset.

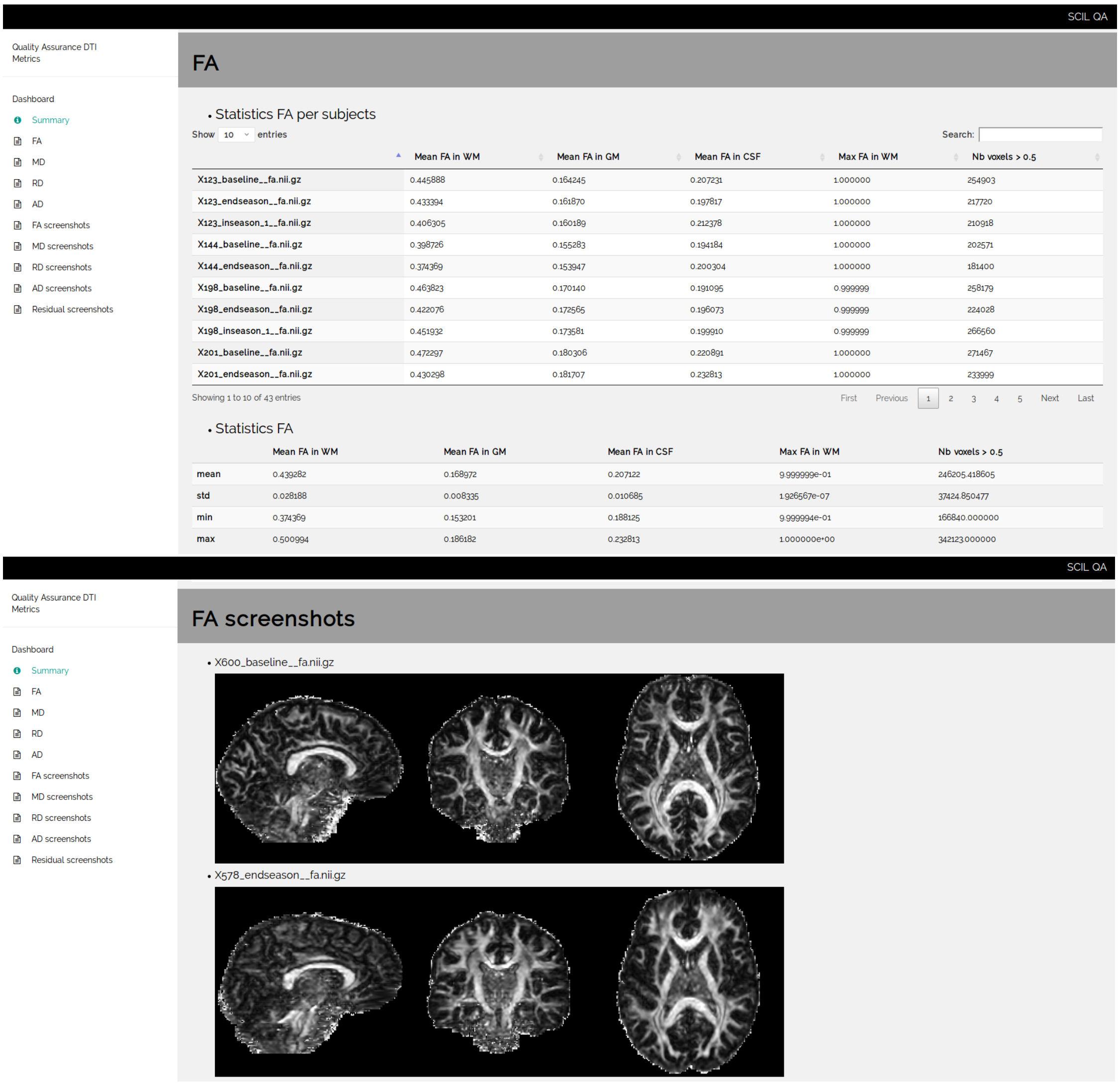


Figure 1: Example of DTI metrics QA/QC report.
